# A Transferable Model For Chromosome Architecture

**DOI:** 10.1101/048439

**Authors:** Michele Di Pierro, Bin Zhang, Erez Lieberman Aiden, Peter G. Wolynes, José N. Onuchic

**Affiliations:** Center for Theoretical Biological Physics, Rice University, Houston, Texas 77005; The Center for Genome Architecture, Baylor College of Medicine, Houston, Texas 77030; Departments of Physics & Astronomy, Chemistry, and Biosciences, Rice University, Houston, Texas 77005

## Abstract

*In vivo*, the human genome folds into a characteristic ensemble of three-dimensional structures. The mechanism driving the folding process remains unknown. We report a theoretical model for chromatin (**Mi**nimal **Chro**matin **M**odel) that explains the folding of interphase chromosomes and generates chromosome conformations consistent with experimental data. The energy landscape of the model was derived by using the maximum entropy principle and relies on two experimentally derived inputs: a classification of loci into chromatin types and a catalog of the positions of chromatin loops. First, we trained our energy function using the Hi-C contact map of chromosome 10 from human GM12878 lymphoblastoid cells. Then we used the model to perform molecular dynamics simulations producing an ensemble of 3D structures for all GM12878 autosomes. Finally, we used these 3D structures to generate contact maps. We found that simulated contact maps closely agree with experimental results for all GM12878 autosomes.

The ensemble of structures resulting from these simulations exhibited unknotted chromosomes, phase separation of chromatin types, and a tendency for open chromatin to lie at the periphery of chromosome territories.

**One Sentence Summary:** We report a model for chromatin that explains and accurately reproduces the three-dimensional structure of chromosomes in interphase.

## Main Text

Chromatin comprises a highly flexible polymer composed of nucleosomes – DNA wrapped around histone proteins – connected to one another by a linker region of 20-50 base pairs (bp). Hundreds of associated structural and regulatory proteins interact with the genetic material coordinating the way chromatin folds to fit inside the nucleus of eukaryotic cells.

The resulting ensemble of partially organized structures brings sections of DNA separated by a great genomic distance into close spatial proximity, and plays an important role in controlling gene transcription (*1, 2*). Although some of the features of this ensemble can be explained using simple polymer physics (*3-5*), there is now ample evidence that specific biochemical interactions play a crucial role (*6-8*). Understanding the interplay between biochemistry, genome architecture, and transcriptional regulation is a major outstanding challenge.

For over two decades, molecular biology techniques that combine chromatin fragmentation and proximity ligation have given us quantitative information about how chromatin is organized *in vivo* (*5, 9-11*). In recent years, Hi-C experiments have made it possible to measure the frequency of contact between all pairs of genomic loci using a single experiment.

Here, we explore a physical model by which local interactions between genomic loci can lead to the conformations of human chromosomes in interphase. Specifically, we propose a theoretical energy landscape model for chromatin folding, designated the **Mi**nimal **Chro**matin **M**odel (MiChroM), which uses the maximum entropy principle (*12, 13*) in combination with a minimal number of assumptions in order to model the structural consequences of the aforementioned biochemical interactions. Importantly, MiChroM can be used to model biochemical interactions even though the identity of the interacting biomolecules is unknown. MiChroM suggests a mechanism that is sufficient to explain chromatin organization and can be used to generate ensembles of 3D structures describing whole genomes. As we will show, contact maps generated *in silico* from these ensembles of structures reproduce in detail the maps from Hi-C.

The first assumption made in MiChroM is that the genome is partitioned into intervals of a handful of types, such that each type of interval is marked by characteristic histone modifications and interacts with a characteristic combination of nuclear proteins. As a result, when two segments of chromatin come into contact, the effective free energy change due to this contact depends, to first order, on the chromatin type of each segment (see also Jost *et al*. (*14))*.

This assumption is supported by both biochemical and structural data. For instance, five distinct types of chromatin have been found in *Drosophila* cells based on the binding patterns of nuclear proteins (*15*). Further, analysis of original Hi-C maps (*5*) suggested that human chromatin is partitioned into two compartments, A and B, each associated with distinct long-range contact patterns. More recently, Rao *et al.* (*8*) used kilobase-resolution Hi-C experiments to show that the human genome can be further partitioned into six subcompartments (A1,2 and B1,2,3,4); each correlated with particular histone marks and associated with a particular pattern of longrange contacts. A similar partitioning of the genome was observed also in mouse (*8, 16*) and *Drosophila* (*17, 18*). Both the boundaries of these genomic intervals and their chromatin types may change along with changes in cell state (*8*). The close association between interval types and long-range contact patterns suggests that intervals of the same type segregate together in the nucleus.

The second assumption made in MiChroM is that certain pairs of genomic “anchor” loci tend to form loops. This tendency is encoded in the model as a change in the effective free energy of a chromatin configuration when the two anchor loci are in contact. This assumption is well-supported by historical literature (*7*), and has been further confirmed by recent high-resolution Hi-C maps of the human genome, where loops are visible as peaks in the contact probability map (*8*). Most loops are associated with convergent pairs of CCCTC binding factor (CTCF)-binding motifs, which have been proposed to help orchestrate loop formation via extrusion (*19*). MiChroM, however, makes no assumption about the particular mechanism of loop formation.

Finally, MiChroM assumes that every time a pair of loci comes into contact there is a gain/loss of effective free energy, *γ*(*d*), that depends only on the genomic distance, *d*. This “ideal chromosome” term models the local structure of chromatin in absence of compartmentalization or looping (*13*), and is sequence translational invariant by construction. The form of the ideal chromosome potential is supported by the widespread evidence that chromatin can behave like a liquid crystal (*20, 21*), and is consistent with the popular notion of the existence of a higher order fiber in chromatin (*22*) while remaining more general.

To build a physical model for chromatin, we use the maximum entropy principle to convert the above three assumptions into an information theoretical energy function. The effective energy that maximizes the information theoretic entropy takes the following form (see SI):

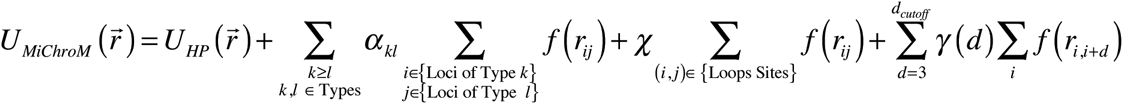

and includes, respectively, the potential energy, 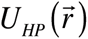, characterizing a generic homopolymer, the interactions between chromatin types (assumption #1), the interactions between loop anchors (assumption #2), and the translational invariant compaction term (assumption #3).

This potential function contains 27 parameters that must be provided in order to fully specify the model. Once the potential function is fully specified, it is possible to perform molecular dynamics simulations of chromatin using as input the classification of loci into chromatin types and the location of loops. This procedure is directly analogous to the simulation of protein folding using amino acid sequence and disulfide bond positions as the only input.

Determining the optimal value for these 27 parameters requires a training data set. In this case, we iteratively adjusted the parameter set in order to reproduce data extracted from a Hi-C contact map of chromosome 10 generated using GM12878 cells (*8*). To do so, we modeled human chromosome 10, which is 136 Mbp long, as a polymer containing 2712 monomers, each representing 50 kb of DNA. We used the annotations generated by Rao *et al.* to assign each monomer a chromatin type, as well as to specify the positions of loops between pairs of monomers. In each iteration, we combined these polymer specifications with the current parameter set in order to generate an ensemble of structures. We then used this ensemble to generate a simulated map of pairwise inter-monomer contact frequencies, and compared this contact map to the one obtained by Rao *et al.* experimentally in order to choose the next set of parameters (see SI).

The simulated contact maps obtained using the final set of parameters correspond closely to the experimental contact maps obtained for chromosome 10 (Pearson’s *r* = 0.95). This correspondence goes beyond the visually obvious “checkerboard” pattern in the simulated contact map (Figure 1). In general, all features larger than 300-400 kb in the experimental contact map (i.e., features that are about an order of magnitude larger than the size of an individual monomer in our simulations) appear to be accurately recapitulated by the MiChroM model. Notably, the power law scaling relationship between the probability of forming contacts and genomic distance, often used to justify the non-equilibrium fractal globule model, is also reproduced with great accuracy by this equilibrium model (Figure 1E).

**Fig. 1.**
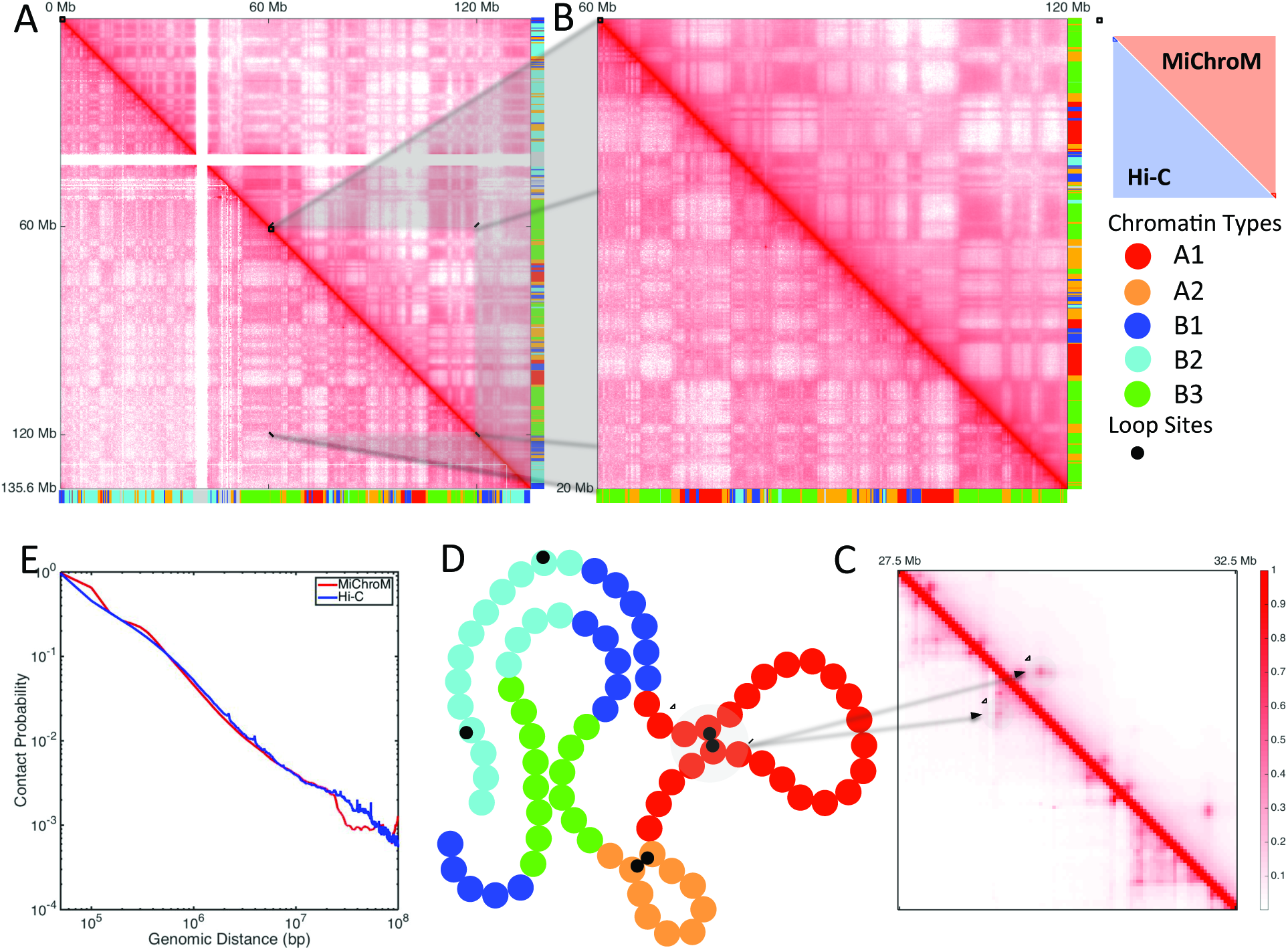
Contact maps obtained by computational modeling using MiChroM closely reproduce experimental maps obtained using the Hi-C protocol. Panels A, B, C show the contact probability map of chromosome 10 of B-lymphoblastoid cells (GM12878). The map obtained from MiChroM is shown in the upper diagonal section of each map while the lower diagonal region shows the maps reported in Rao *et al* (*8*). A symmetric figure indicates that the experimental data are well reproduced. Color bars on axes show the chromatin type sequences of chromosome 10. A) Complete contact map of chromosome 10 (log scale). B) Magnification of the 60-120 Mb region of chromosome 10 (log scale). At this magnification the relationship between chromatin types and spatial proximity is clearly visible. MiChroM accurately reproduces the pattern of contacts measured by Hi-C. C) Further magnification of the 27.5-32.5 Mb region (linear scale). The peaks in contact probability characterizing the loops are clearly visible in both experimental and computational maps (See SI for more detail). D) Schematic representation of MiChroM. Each bead represents 50 kb of chromatin belonging to one single chromatin type represented by its specific color as in color bars of panels A and B. A smaller black bead marks the location of loop sites. E) The probability of contacts as a function of genomic distance from experiment and *in silico.*

Next, we applied the MiChroM model to the remaining GM12878 autosomes by combining the potential function with the experimentally derived monomer type and loop annotations. When each chromosome is simulated separately, the resulting intrachromosomal contact map closely corresponds to the experimental contact map in every case. Notably, the correspondence for autosomes that were not used to train the potential function was typically as close (Pearson’s *r* = 0.95) as the correspondence for chromosome 10 (See Figure 2, S2-S47, and SI).

**Fig. 2.**
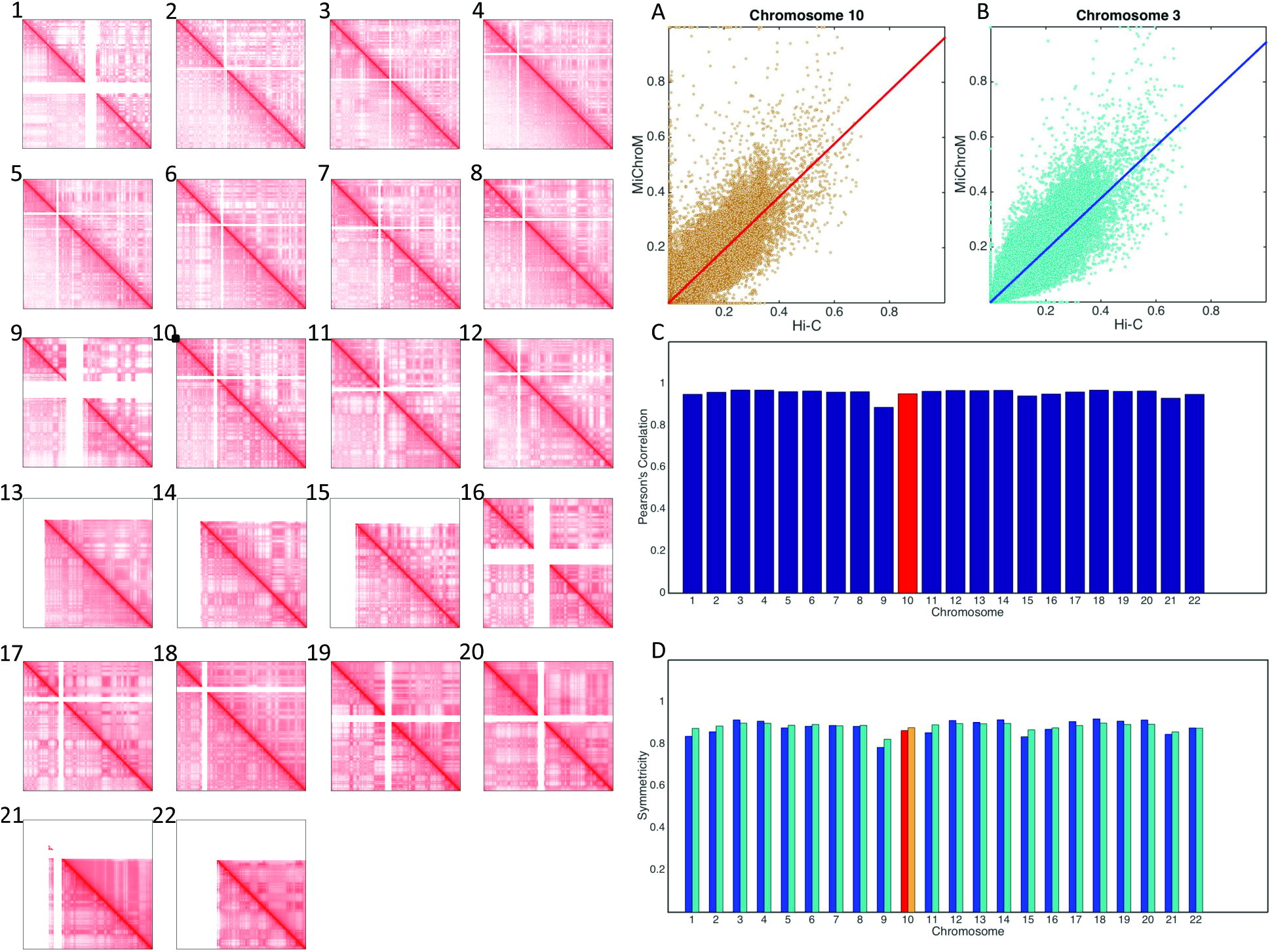
MiChroM generates 3D structures for the chromosomes of B-lymphoblastoid cells (GM12878) that are in agreement with experimental data. Panels 1-22 show contact maps of chromosomes 1-22 represented in log scale; upper diagonal regions show results from computational modeling and lower diagonal regions show maps obtained using Hi-C protocol (*5*). Chromosome 10 was the only chromosome used for calibration. The contact maps for the remaining 21 chromosomes were generated *de novo* using the chromatin type sequence and location of loops as only inputs. The quality of the generated contact maps is consistently high, as shown by the symmetry of the maps. Panels A, B, C, and D show different measures of the accuracy of the maps generated by our model. A, B) Scatter plots of Hi-C vs MiChroM contact probability for all the possible contacts together with a linear fit of the data obtained by using the least squares method. Panel A shows data from chromosome 10, the slope of the fitting function is 0.96 and the intercept is 0.0003. Panel B shows data from chromosome 3, the slope in this case is 0.94 and the intercept is 0.0003. The experimental and computational data sets show a strong linear relationship confirming their similarity. The quality of such linear relationship rema<u>ins</u> substantially unaltered whether we look at chromosome 10, which was used for calibration, or at chromosome 3 that was independently generated. C) Pearson’s correlation between the contact probabilities generated by MiChroM and as measured by Hi-C for all the 22 chromosomes. The average correlation is 0.956. For chromosome 10, which was used for calibration, the correlation is 0.952 (See also figure S2-S3). D) Symmetry score (See SI for definition) for the combined Hi-C/MiChroM maps for the 22 chromosomes. The score is bounded by 0 and 1, with 1 indicating a perfectly symmetric matrix (i.e. a perfect agreement between model and experiments). The average symmetry score is 0.88 using both the 2-norm (blue bars) and the Frobenius norm (cyan).

When we examined the ensemble of 3D structures for each individual chromosome, we observed that each chromosome formed a compact chromosome territory. We also observed the phase separation of chromatin types within this territory, leading to subvolumes comprising only a single type of genomic interval (Figure 3A). Usually, only a single subvolume formed for each subcompartment, although in some cases we observed multiple subvolumes of a single type. Similarly, we see that highly expressed genes (as measured by RNA-Seq (*23*)) tend to occupy spatial subvolumes, which is expected given that highly expressed genes lie predominantly in the A compartment. Overall, these findings are consistent with the notion that different types of intervals co-localize in distinct spatial compartments. Interestingly, the A compartment tends to be less densely packed and to lie at the periphery of the chromosome territory. These observations are consistent with the findings of prior studies using both microscopy and Hi-C (*8, 24, 25*). Notably, a control model composed of a simple self-avoiding homopolymer chain failed to exhibit any of these results, and instead recapitulated the expected properties for an equilibrium globule (Figures 3A and 3B).

**Fig. 3.**
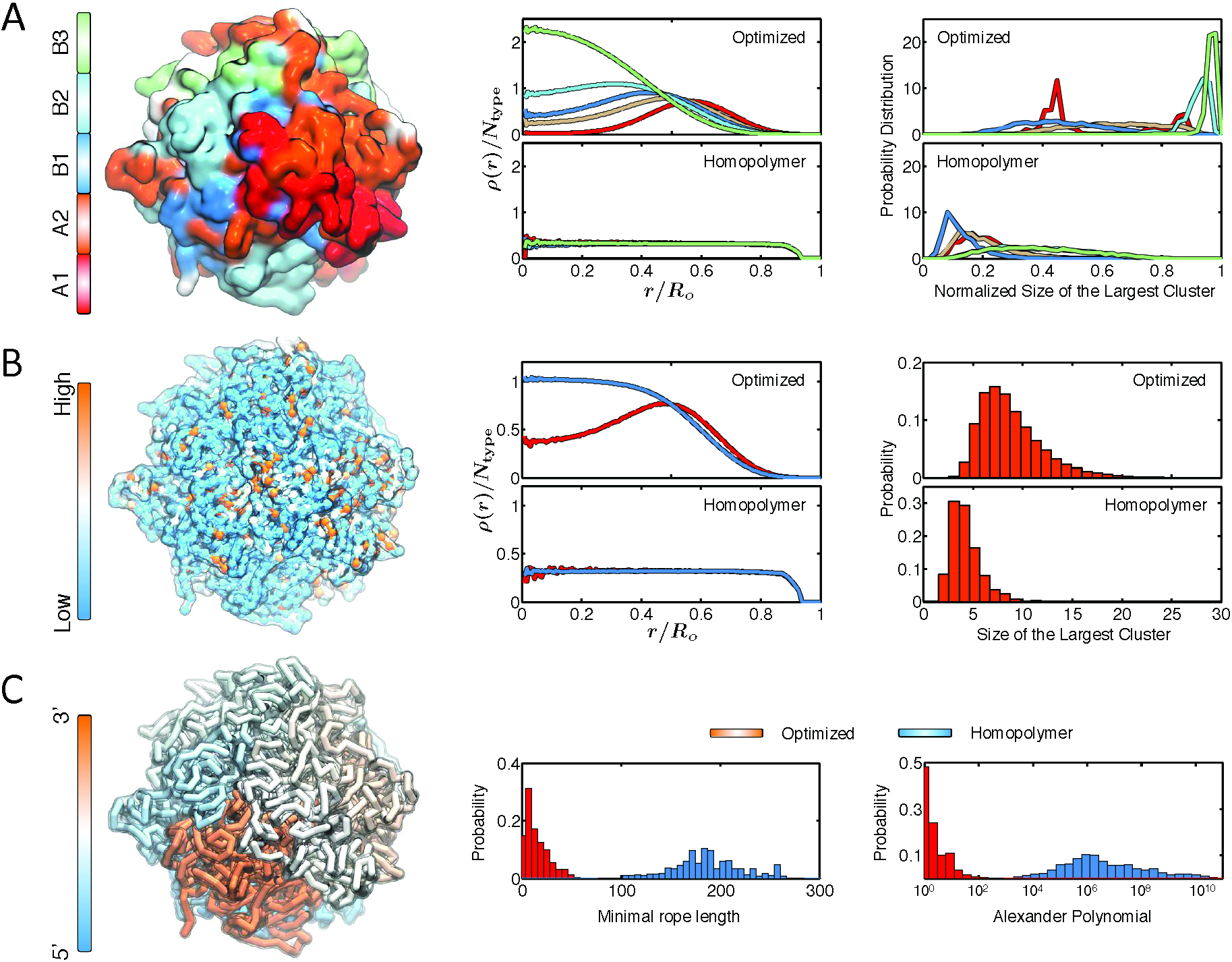
Structural characterization of the simulated conformations for chromosome A) Chromatin of different types phase separate, with A types localizing at the surface and B types in the interior. (*Left*) Surface plot for the chromosome colored by chromatin types, with the coloring scheme shown at the side. (*Middle*) Radial density profiles for different chromatin types calculated from the ensemble of simulated chromosome (top) and homopolymer (bottom) conformations. (*Right*) Probability distributions for the size of the largest cluster found for each chromatin type from the ensemble of simulated chromosome (top) and homopolymer (bottom) conformations. The cluster sizes for different types are normalized by the total number of genomic loci for that type. B) Genomic loci with high gene expression spatially colocalize at the exterior of the chromosome. (*Left*) A chromosome structure colored by gene expression with red and blue representing high and low expression, respectively. (*Middle*) Radial density profiles for genomic loci with high and low gene expression calculated from the ensemble of simulated chromosome (top) and homopolymer (bottom) conformations. (*Right*) Probabilities for finding the largest cluster of size *N* for highly expressed genomic loci from the ensemble of simulated chromosome (top) and homopolymer (bottom) conformations. C) Simulated chromosomes adopt knot-free conformations. (*Left*) A chromosome structure colored by genomic distances, with one end of the chromosome shown in blue and the other end in red. (*Middle*) Probability distributions of the knot invariant measured as minimal rope length for the ensemble of simulated chromosome (red) and homopolymer (blue) conformations. (*Right*) Probability distributions of the knot invariant measured as Alexander polynomial for the ensemble of simulated chromosome (red) and homopolymer (blue) conformations.

It is commonly assumed that one essential feature of chromosomes is the absence of knots, as one might suppose that a highly knotted structure could create obstacles to the transcription process. We studied the extent of knotting in the ensemble of chromosome structures sampled from the optimized energy landscape and from the homopolymer potential. In order to quantify knotting in a particular conformation of the chromosome, we used two different knot invariants: the Alexander polynomial and the minimal rope length required to generate a topologically equivalent knot (*13, 26*). Both measures show that the configurations produced by MiChroM are largely devoid of knots. In contrast, the homopolymer control system tended to form extraordinarily complex knots (Figure 3C). This topological feature is a direct result of inferring the energy landscape from the three physical assumptions explained above. Remarkably, the simple equilibrium mechanism underlying MiChroM produces ensembles of structures that are devoid of knots.

Finally, we used MiChroM to jointly simulate chromosomes 17 and 18 (Figure S1). This allowed us to explore whether the MiChroM potential function, which was trained using a single intrachromosomal contact map for chromosome 10, could successfully reproduce genome architecture at a larger scale. The resulting intrachromosomal contact maps are essentially the same as those simulated in isolation (Pearson’s *r* = 0.96). The phenomenon of phase separation of chromatin types now extends to both chromosomes creating larger regions of space occupied by one single type. Spatial confinement introduces artifacts in the frequency of interchromosomal contacts; therefore, the interchromosomal contact map from simulation shows somewhat increased probabilities with respect to Hi-C. Even with the biased intensity, the two-chromosome map shows a correct pattern of interchromosomal interactions.

When we examined the 3D ensemble, we found that, despite the extensive contacts between the chromosomes, the chromosomes were not entangled with one another (Figure S1B); instead, we observed the formation of non-overlapping chromosome territories. This last result highlights the fact that MiChroM can successfully recapitulate features of the nucleus as a whole.

The **Mi**nimal **Chro**matin **M**odel assumes that chromosomes fold under the action of a cloud of proteins that bind with different selectivity to different sections of chromatin, and offers a simple strategy for recapitulating the energy landscape created by such interactions. This energy landscape brings about transient contacts rather than permanent ones, which is consistent with the fact that most of the experimentally observed contacts between two genetic loci only occur in a small fraction of cells at a given time (*5, 27*). Contacts associated with loop formation tend to be more frequent; accordingly, our optimization algorithm assigns them a larger free energy gain upon formation. In humans, we find that six types of chromatin are sufficient to reproduce the arrangement of interphase DNA *in vivo.* The fact that our model can be reliably transferred from one chromosome to the rest suggests the plausibility of the proposed energetic mechanism, even if the underlying biochemical details remain unclear at the present time.

As shown, MiChroM is able to explain and reproduce the results of DNA proximity ligation experiments. Nevertheless, caution must be applied in the interpretation of these results. Hi-C experiments are performed using millions of cells at once, and report only a population average. We know little about what happens in individual cells at specific moments in time. For instance, a typical cell population interrogated by Hi-C may contain entirely separate subpopulations, as well as fluctuating or even oscillating configurations. These would be lost in MiChroM.

The classification of loci into chromatin types and the position of chromatin loops, which are inputs of our model, are strongly associated with epigenetic features (histone modifications and bound CTCF motifs in convergent orientation) that can be directly and inexpensively assayed by ChIP-Seq. Exploiting these associations along with MiChroM opens up the possibility of predicting *in silico* the 3D structure of whole genomes starting from 1-dimensional genomics data, which are often already publicly available.

## Acknowledgments

We thank Ryan R. Cheng, Davit Potoyan and Lena Simine for many useful discussions, and Erica J. Di Pierro for help in editing the manuscript.

This work was supported by the Center for Theoretical Biological Physics sponsored by the National Science Foundation (grants PHY-1427654 and NSF-MCB-1214457) and by the Cancer Prevention and Research Institute of Texas (CPRIT – grant R1110). Michele Di Pierro was also supported by the Welch Foundation (grant C-1792).

## References

1. T. Cremer, C. Cremer, Chromosome territories, nuclear architecture and gene regulation in mammalian cells. Nat Rev Genet 2, 292–301 (2001).

2. W. A. Bickmore, The Spatial Organization of the Human Genome. Annu Rev Genom Hum G 14, 67–84 (2013).

3. A. Y. Grosberg, S. K. Nechaev, E. I. Shakhnovich, The Role of Topological Constraints in the Kinetics of Collapse of Macromolecules. JPhys-Paris 49, 2095–2100 (1988).

4. A. Grosberg, Y. Rabin, S. Havlin, A. Neer, Crumpled Globule Model of the 3-Dimensional Structure of DNA. Europhys Lett 23, 373–378 (1993).

5. E. Lieberman-Aiden et al., Comprehensive Mapping of Long-Range Interactions Reveals Folding Principles of the Human Genome. Science 326, 289–293 (2009).

6. B. van Steensel, Chromatin: constructing the big picture. EMBO J 30, 1885–1895 (2011).

7. J. E. Phillips, V. G. Corces, CTCF: Master Weaver of the Genome. Cell 137, 1194–1211 (2009).

8. S. S. P. Rao et al., A 3D Map of the Human Genome at Kilobase Resolution Reveals Principles of Chromatin Looping. Cell 159, 1665–1680 (2014).

9. K. E. Cullen, M. P. Kladde, M. A. Seyfred, Interaction between Transcription Regulatory Regions of Prolactin Chromatin. Science 261, 203–206 (1993).

10. J. Dekker, K. Rippe, M. Dekker, N. Kleckner, Capturing chromosome conformation. Science 295, 1306–1311 (2002).

11. J. Dostie et al., Chromosome Conformation Capture Carbon Copy (5C): A massively parallel solution for mapping interactions between genomic elements. Genome Res 16, 1299–1309 (2006).

12. E. T. Jaynes, Information Theory and Statistical Mechanics. Phys Rev 106, 620–630 (1957).

13. B. Zhang, P. G. Wolynes, Topology, structures, and energy landscapes of human chromosomes. P Natl Acad Sci USA 112, 6062–6067 (2015).

14. D. Jost, P. Carrivain, G. Cavalli, C. Vaillant, Modeling epigenome folding: formation and dynamics of topologically associated chromatin domains. Nucleic Acids Res 42, 9553–9561 (2014).

15. G. J. Filion et al., Systematic protein location mapping reveals five principal chromatin types in Drosophila cells. Cell 143, 212–224 (2010).

16. J. R. Dixon et al., Topological domains in mammalian genomes identified by analysis of chromatin interactions. Nature 485, 376–380 (2012).

17. K. P. Eagen, T. A. Hartl, R. D. Kornberg, Stable Chromosome Condensation Revealed by Chromosome Conformation Capture. Cell 163, 934–946 (2015).

18. T. Sexton et al., Three-Dimensional Folding and Functional Organization Principles of the Drosophila Genome. Cell 148, 458–472 (2012).

19. A. L. Sanborn et al., Chromatin extrusion explains key features of loop and domain formation in wild-type and engineered genomes. P Natl Acad Sci USA 112, E6456–E6465 (2015).

20. E. Boy de la Tour, U. K. Laemmli, The metaphase scaffold is helically folded: sister chromatids have predominantly opposite helical handedness. Cell 55, 937–944 (1988).

21. N. Naumova et al., Organization of the Mitotic Chromosome. Science 342, 948–953 (2013).

22. K. Maeshima, S. Hihara, M. Eltsov, Chromatin structure: does the 30-nm fibre exist in vivo? Curr Opin Cell Biol 22, 291–297 (2010).

23. A. Mortazavi, B. A. Williams, K. Mccue, L. Schaeffer, B. Wold, Mapping and quantifying mammalian transcriptomes by RNA-Seq. Nat Methods 5, 621–628 (2008).

24. A. N. Boettiger et al., Super-resolution imaging reveals distinct chromatin folding for different epigenetic states. Nature 529, 418-+ (2016).

25. M. R. Hubner, D. L. Spector, Chromatin Dynamics. Annu Rev Biophys 39, 471–489 (2010).

26. A. Stasiak, V. Katritch, L. H. Kauffman, Ideal knots. K & E series on knots and everything (World Scientific, Singapore; River Edge, NJ, 1998), pp. x, 414 p.

27. F. Bantignies et al., Polycomb-Dependent Regulatory Contacts between Distant Hox Loci in Drosophila. Cell 144, 214–226 (2011).

